# Optimized detection of allelic imbalances specific for homologous recombination deficiency improves the prediction of clinical outcomes in cancer

**DOI:** 10.1101/2021.08.19.456809

**Authors:** Fernando Perez-Villatoro, Jaana Oikkonen, Julia Casado, Anastasiya Chernenko, Doga C. Gulhan, Manuela Tumiati, Yilin Li, Kari Lavikka, Sakari Hietanen, Johanna Hynninen, Ulla-Maija Haltia, Jaakko S. Tyrmi, Hannele Laivuori, Panagiotis A. Konstantinopoulos, Sampsa Hautaniemi, Liisa Kauppi, Anniina Färkkilä

## Abstract

Homologous recombination DNA-repair deficiency (HRD) is a common driver of genomic instability and confers a therapeutic vulnerability in cancer. The accurate detection of somatic allelic imbalances (AIs) has been limited by methods focused on *BRCA1/2* mutations and using mixtures of cancer types. Using pan-cancer data, we revealed distinct patterns of AIs in high-grade serous ovarian cancer (HGSC). We used machine learning and statistics to generate improved criteria to identify HRD in HGSC (ovaHRDscar). ovaHRDscar significantly predicted clinical outcomes in three independent patient cohorts with higher precision than previous methods. Characterization of 98 spatiotemporally distinct metastatic samples revealed low intra-patient variation and indicated the primary tumor as the preferred site for clinical sampling in HGSC. Further, our approach improved the prediction of clinical outcomes in triple-negative breast cancer (tnbcHRDscar), validated in two independent patient cohorts. In conclusion, our tumor-specific, systematic approach has the potential to improve patient selection for HR-targeted therapies.

## BACKGROUND

As a part of the Fanconi Anemia (FA) pathway, homologous recombination (HR) is an evolutionarily conserved, tightly regulated mechanism for high-fidelity repair of DNA double-strand breaks (DSBs)^1^. Deficiency in homologous recombination (HRD) has profound consequences for replicating cells driving genomic instability and oncogenic transformation. In cancer, HRD results in a fundamental vulnerability, and tumors with HRD are markedly sensitive to DSB-inducing agents such as platinum-based chemotherapy and Poly-ADP Ribose Polymerase (PARP) inhibitors^2^.

High-grade serous ovarian cancer (HGSC), the most common and most lethal subtype of ovarian cancers^3^, is characterized by profound genomic instability. Around half of the HGSC cases harbor genomic alterations leading to HRD^4^, and these patients have been shown to benefit from treatment with PARP inhibitors ^5,6^. The HRD test previously used in PARP inhibitor clinical trials (MyriadMyChoise®CDx)^5,6^ works by quantifying specific allelic imbalances (AIs): 1) Large scale transitions (LSTs)^7^, 2) Loss of heterozygosity (LOH)^8^ and 3) Telomeric allelic imbalances (TAIs)^9^. However, the decision criteria for these HRD-specific AIs (HRD-AIs) and the HRD status classification were originally designed using a mixture of breast and ovarian cancer samples^7,8,9,10^. Further, other algorithms for HRD detection have primarily focused on *BRCA1/2* mutation prediction^11,12^. As the genomic drivers and mutational processes differ across the cancer types, the details of the genomic instability occurring due to HRD in HGSC remain unclear.

Herein, via pan-cancer analysis, we show that HGSC harbors unique patterns of AIs, which are also distinct from triple-negative breast cancers (TNBC). Using a systematic approach based on machine learning and statistics on The Cancer Genome Atlas ovarian cancer (OVA-TCGA) multi-omics dataset, we optimized the criteria for HRD-AIs on HGSC. We implemented these criteria as an open-source algorithm (ovaHRDscar) to reliably define HRD status beyond the prediction of *BRCA1/2* mutations. We show that ovaHRDscar improves the prediction of clinical outcomes in three independent clinical datasets compared to previous algorithms. Further, we show that our approach improves the prediction of clinical outcomes also in TNBC (tnbcHRDscar). Thus, our machine learning-aided disease-specific approach (HRDscar) shows promise as a biomarker that can improve outcome prediction and patient selection for HR-targeted therapies in cancer.

## RESULTS

### Systematic pan-cancer characterization reveals unique features of allelic imbalances in HGSC

To elucidate the potential differences in the patterns of AIs across human cancers, we first characterized the quantity and the length distributions of AIs in the 18 most common cancer types from the TCGA (**Fig. 1a**). Interestingly, HGSC had the highest number of AIs (**Fig. 1b**) and the lowest median length (**Fig. 1c**). Concordantly, HGSC showed the highest levels of LOH events (**Sup. Fig. 1a**) with one of the lowest median length (**Sup. Fig. 1b**).

**Figure 1.**
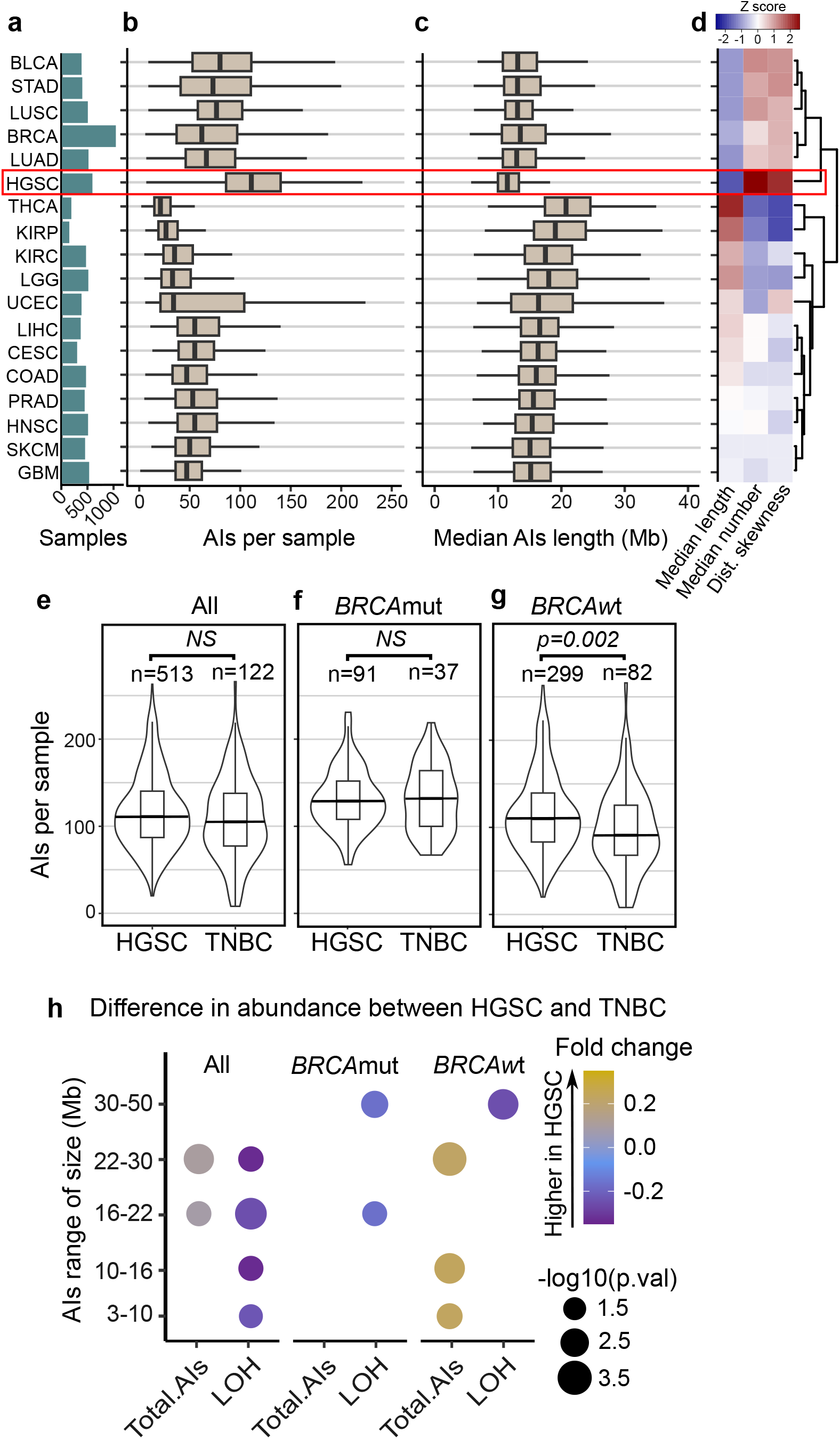
Pan-cancer characterization of AIs reveals unique patterns in HGSC. **a** Types of cancer with more than 200 samples in TCGA and their corresponding number of samples are shown in green bars; bladder urothelial carcinoma (BLCA), stomach adenocarcinoma (STAD), lung squamous cell carcinoma (LUSC), breast invasive carcinoma (BRCA), thyroid carcinoma (THCA), kidney renal papillary cell carcinoma (KIRP), kidney renal clear cell carcinoma (KIRC), brain Lower Grade Glioma (LGG), uterine Corpus endometrial carcinoma (UCEC), liver hepatocellular carcinoma (LIHC), cervical squamous cell carcinoma and endocervical adenocarcinoma (CESC), colon adenocarcinoma (COAD), prostate adenocarcinoma (PRAD), head and neck squamous cell carcinoma (HNSC), skin cutaneous melanoma (SKCM), glioblastoma multiforme (GBM),. **b** Box plots representing the number of AIs longer than 3Mb and smaller than 50Mb per sample. HGSC showed the highest average levels of AIs. **c** Box plots showing the median length of AIs (longer than 3Mb and smaller than 50Mb) per sample. HGSC showed the lowest median length of AIs per sample. **d** Hierarchical clustering for the types of cancer using as variables the median length, the median number of AIs per sample, and the skewness of the distribution of AIs length. **e** Violin- and box plots representing the number of AIs per sample. A long vertical line represents the median, HGSC showing a similar number of AIs as compared to TNBC (U test). **f** Comparison of *BRCA*mut samples showing similar abundances of AIs in HGSC as compared to TNBC (U test). **g** The *BRCA*wt samples showing significantly higher number of AIs in HGSC than in TNBC (U test, p=0.002). **h** Dot plot showing the difference in abundance for AIs of specific length between HGSC and TNBC. The dot sizes represent the p-values (U-test) and dot colors represent the fold-change (ratio of HGSC/TNBC abundance of AIs minus one), only dots for corresponding significant differences are shown (U test, p < 0.05).

We next performed hierarchical clustering using the median length and number of AIs per sample and the skewness of the length distribution of the AIs for each cancer type. This analysis shows two main clusters: the first cluster consisting of six cancer types (bladder urothelial carcinoma (BLCA), stomach adenocarcinoma (STAD), lung squamous cell carcinoma (LUSC), lung adenocarcinoma (LUAD), breast invasive carcinoma (BRCA), and HGSC) with a higher amount but a lower median length of AIs (upper cluster: **Fig. 1d**). The second cluster consisting of the remaining 12 cancer types (lower cluster: **Fig. 1d**). The same main clusters were observed when using only LOH events (**Sup. Fig. 1c**).

As TNBC and HGSC are enriched in *BRCA1/2* genetic mutations (*BRCA*mut)^13^, both cancers were used to define the HRD-algorithm in the MyriadMyChoise®CDx assay by Telli et al.^10^. We next compared the differences in AIs between these two cancer types. We observed a significant difference in the abundance of AIs between HGSC and TNBC, specifically among the *BRCA1/2-*wild-type (*BRCAwt*) tumors (U test, p = 0.002, **Fig. 1e to g**). Interestingly, HGSC had lower levels of LOH events than TNBC (U test, p = 0.002, **Sup. Fig. 1d**), also among the *BRCA*mut samples (U test, p = 0.049, **Sup. Fig. 1e**) but not in the *BRCA*wt samples (**Sup. Fig. 1f**). Overall, HGSC showed a higher number of AIs of different lengths, while TNBC had a higher number of LOH events (**Fig. 1h**). These results highlight the distinct characteristics of AI events in HGSC, especially among the *BRCA*wt tumors, compared to other cancer types.

### Machine learning-aided detection of HRD-specific AIs improves the detection of HRD in HGSC

Although a wide range of molecular alterations is known to cause HRD, previous studies have focused on *BRCA1/2* mutations to detect HRD-specific AIs (HRD-AIs), potentially failing to detect non-*BRCA* associated HRD alterations while losing specificity to classify the HR-proficient (HRP) samples accurately. To this end, we aimed to identify AIs overrepresented in samples carrying a wider range of genetic alterations (mutations, gene deletions, promoter hypermethylation) associated with HRD in HGSC (**Fig. 2a**). To generate accurate selection criteria for HRD-AIs, we utilized SNP-arrays data from HGSC samples from TCGA (OVA-TCGA) and its associated genomic and DNA methylation data. Using prior knowledge and multi-omics data, we annotated 115 HRD samples harboring a somatic or germline mutation, gene deletion, or promoter hypermethylation in the *BRCA1/2* or *RAD51* paralog genes, and 29 HRP samples that did not harbor any of the alterations used to select the HRD samples, nor deletions in any other HR-related gene (**Fig. 2a**). A detailed description of the genomic alterations in the samples is reported in **Sup. Table 1**. Overall, the HRD samples had a higher number of all AIs than the HRP samples (U test, p=0.0028, **Sup. Fig. 2a**). Importantly, HRD samples had a notably higher proportion of AIs of a specific length that spanned from 1Mb to 30Mbs. In contrast, the HRP samples contained a higher proportion of AIs and LOH events smaller than 1Mb (**Sup. Figs. 2b, 2c**).

**Figure 2.**
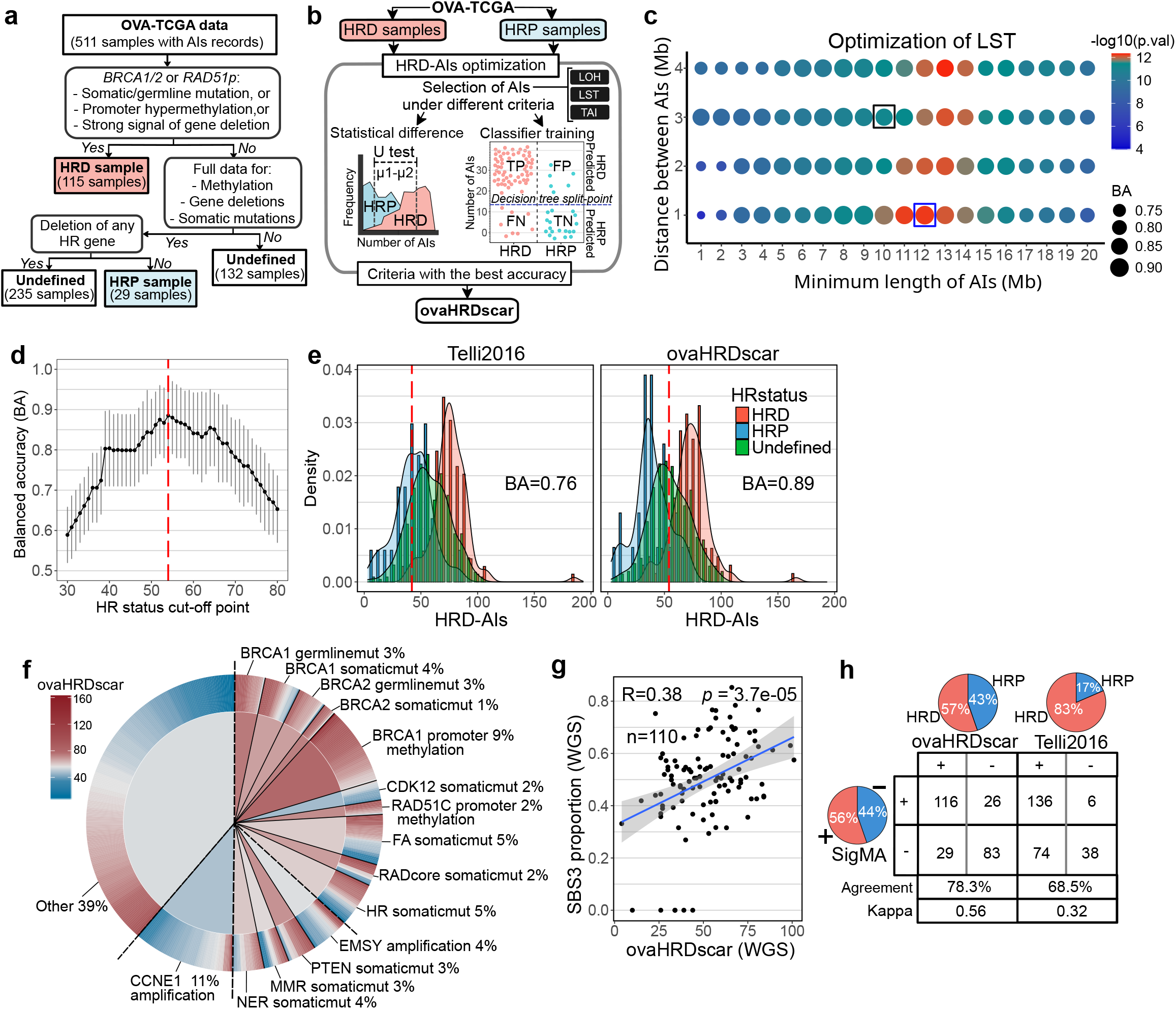
Machine learning-aided detection of AIs associated with HRD shows improved accuracy and correlations with genomic features of HRD in HGSC. **a** Selection criteria for annotating HRD, HRP and undefined HGSC samples in the OVA-TCGA. **b** A scheme of the approach used to generate accurate criteria for selecting HRD-AIs in HGSC samples. **c** For LST events, the size of dots represents the decision tree balanced accuracy (BA) of classifying HRD and HRP when selecting AIs of the corresponding criteria, the dot colors represent the statistical difference (U test, p-value) in abundance of AIs between HRD and HRP samples. The black box corresponds to the selection criteria proposed by Telli2016, the blue box correspond to the best BA and U-test value. **d** Evaluation of ovaHRDscar cut-off to define HR-status. The black dots connected with a line correspond to the balanced accuracy (BA) of the classification of the annotated HRD and HRP samples using the given cut-off value, the 95% confidence intervals are shown in grey vertical lines, value of 54 (red dashed line) corresponds to the highest BA. **e** Density distribution of HRD-AIs according to Telli2016 and ovaHRDscar algorithms. The red dashed line represents the cut-off established to define the HR-status using Telli2016 (≥ 42) and using ovaHRDscar (≥ 54). The BA of classification of the annotated HRD and HRP is shown, density distribution colors correspond to the samples annotated as in the panel a. **f** Levels of ovaHRDscar in OVA-TCGA samples harboring different genetic or epigenetic alterations associated with HRD in HGSC^4^. The colors correspond to the ovaHRDscar; in the outer ring of the pie chart every line represents a sample and in the center of the pie chart the colors correspond to the average number of HRD-AIs per genetic or epigenetic alteration. For the somatic mutations (somaticmut) gene deletions were included. **g** Linear regression of the proportion of single base substitution signature 3 (SBS3) and the ovaHRDscar levels in PCAWG samples (Pearson r’=0.38). Blue line shows the regression line and the 95% confidence intervals are shown in grey. **h** The SBS3 status inferred using SigMA^16^ showing a higher agreement with ovaHRDscar (agreement=78.3%, Cohen’s kappa = 0.56) than with the Telli2016 algorithm (agreement=68.5%, Cohen’s kappa = 0.32). In the pie charts and table + and - correspond to the number of HRD positive and HRD negative samples identified under each criterion, respectively. On the bottom is shown the number of samples and the level of agreement between the corresponding criteria.

We next applied statistics and machine learning^14^ to identify the specific length and selection criteria of LOH, LST, and TAI events overrepresented in the HRD samples (**Fig. 2b**). We then compared the accuracies of the herein optimized criteria for HRD to those used in Telli et al.^10^ (hereafter called Telli2016). Notably, for LSTs, our approach increased the accuracy of classification of the HRD/HRP samples from 86% to 90% when using the new criteria (**Fig. 2c**). For LOH events, the accuracy increased from 85% to 88% when using the new criteria (**Sup. Fig. 2d**). We also assessed the HRD classification accuracy of LSTs consisting of three consecutive AIs. However, this produced a lower accuracy (**Sup. Fig. 2e**). The largest improvement in accuracy occurred after including all TAIs larger than 1Mb, and the accuracy for HRD-specific TAI events increased from 67% to 78% when compared to the Telli2016 criteria (**Sup. Fig. 2f).**

Via our systematic approach, we observed the following AIs to be most characteristic of HRD in HGSC: 1) LOH > 15Mb and <50Mb, 2) for LSTs AI > 12Mb, with a distance between them <1Mb, and 3) TAI >1Mb. The sum of these events is hereafter called the ovaHRDscar levels. Then, using bootstrapping subsampling of the pre-annotated HRD and HRP samples, we evaluated the optimal cut-off value for ovaHRDscar to define the final HR-status as HRD or HRP. The value with the highest balanced accuracy (BA) was 54 (**Fig. 2d**), meaning that values higher or equal than 54 correspond to HRD, with higher accuracy for HR-status classification (BA=0.89, right panel **Fig. 2e**) as compared to the Telli2016 algorithm (BA=0.76, left panel **Fig. 2e**). In addition, using a HRD/HRP cut-off value of 54 in the Telli2016 algorithm (hereafter Telli2016-54), the BA remained below that of ovaHRDscar (0.86 vs 0.89, **Sup. Fig. 2g**).

### ovaHRDscar levels correlate with genomic features of HRD and show concordance in WGS data

To investigate the relationships of ovaHRDscar with other known genomic features associated with HRD, we annotated the OVA-TCGA samples according to mutations, gene deletions, and promoter hypermethylation patterns previously reported to be associated with HRD^4^ (**Fig. 2f**). On average, samples with somatic mutations in *BRCA1, BRCA2, PTEN*, or somatic mutations or gene deletions in any gene belonging to the Fanconi Anemia (FA) or HR pathways showed high ovaHRDscar levels. Likewise, samples that contained hypermethylation in the promoter regions of *BRCA1* or *RAD51C* genes or germline mutations in *BRCA1* or *BRCA2* had, on average, high ovaHRDscar levels. As expected, samples harboring an amplification in *CCNE1* (**Sup. Fig. 2h**) had significantly lower levels of ovaHRDscar. However, samples with *EMSY* amplification and *CDK12* somatic mutation did not result in higher ovaHRDscar levels than CCNE1 amplified samples (**Sup. Fig. 2h**).

To assess the concordance of ovaHRDscar between SNP array and whole genome sequencing (WGS) data, we next quantified the ovaHRDscar levels in HGSC samples from the Pan-Cancer Analysis of Whole Genomes project (PCAWG)^15^. The ovaHRDscar levels were highly concordant between WGS and SNP-arrays (Lin’s concordance correlation coefficient, ccc = 0.90; **Sup. Fig. 2i**) in 41 OVA-TCGA samples that were also included in the PCAWG project, consistent with a previous report in breast cancer samples^16^. Next, we tested the correlation of ovaHRDscar with the single base substitution signature 3 (SBS3), which has been associated with HRD^17^. We found that the ovaHRDscar levels detected in WGS positively correlated with the proportion of SBS3 in WGS (Pearson, r’=0.38, p= 3.7e-05; **Fig. 2g**). The SBS3 proportions also correlated with the number of HRD-AIs using the Telli2016 algorithm in the PCAWG cohort (**Sup. Fig. 2j**). We next compared the performance of ovaHRDscar to that of SBS3 inferred from whole exome sequencing (WES) data with a likelihood-based approach SigMA^18^, in 254 samples from the OVA-TCGA. The ovaHRDscar algorithm detected 57% of samples as HRD, and the SigMA tool classified 56% of samples as SBS3+; in contrast, the Telli2016 algorithm identified 83% of the samples as HRD (**Fig. 2h**). HRD detection with ovaHRDscar showed a higher agreement with SigMA (agreement 78.3% and Cohen’s kappa = 0.56) as compared to the Telli2016 algorithm (agreement 68.5% and Cohen’s kappa = 0.32; **Fig. 2h**) or to the Telli2016-54 (agreement 77.2% and Cohen’s kappa = 0.53; **Sup. Fig. 2k**).

### ovaHRDscar improves the prediction of PFS and OS compared to previous algorithms

Next, we measured the association of HR-status classification by ovaHRDscar to progression-free survival (PFS, see methods) in advanced HGSC patients treated with platinum-based chemotherapy in the TCGA and an independent prospective validation dataset (HERCULES). We compared the performance of the ovaHRDscar to *BRCA1/2* deficiency status to the Telli2016 algorithm. The Telli2016 algorithm uses a cut-off value of 63, as proposed by Takaya et al.^19^. As *BRCA1/2* mutations can affect patient outcomes, we assessed the performances of ovaHRDscar in the TCGA dataset after excluding the samples used when defining ovaHRDscar, even though clinical outcomes were not utilized for designing the criteria of ovaHRDscar. *BRCA1/2* mutation or deletion status (*BRCA*mut/del) was not significantly associated with PFS (Log-rank p=0.72; **Fig. 3a**). For OVA-TCGA (**Fig. 3a to 3c**), we found that ovaHRDscar positivity was associated with prolonged PFS (Log-rank p=4.4e-04; **Fig. 3c**). Consistently, ovaHRDscar positive patients had a longer PFS in the independent HERCULES validation cohort (Log-rank p=0.001; **Sup. Fig. 3a to 3c**), while the Telli2016 algorithm did not reach statistical significance in predicting PFS (Log-rank p=0.11; **Sup. Fig. 3b**).

**Figure 3.**
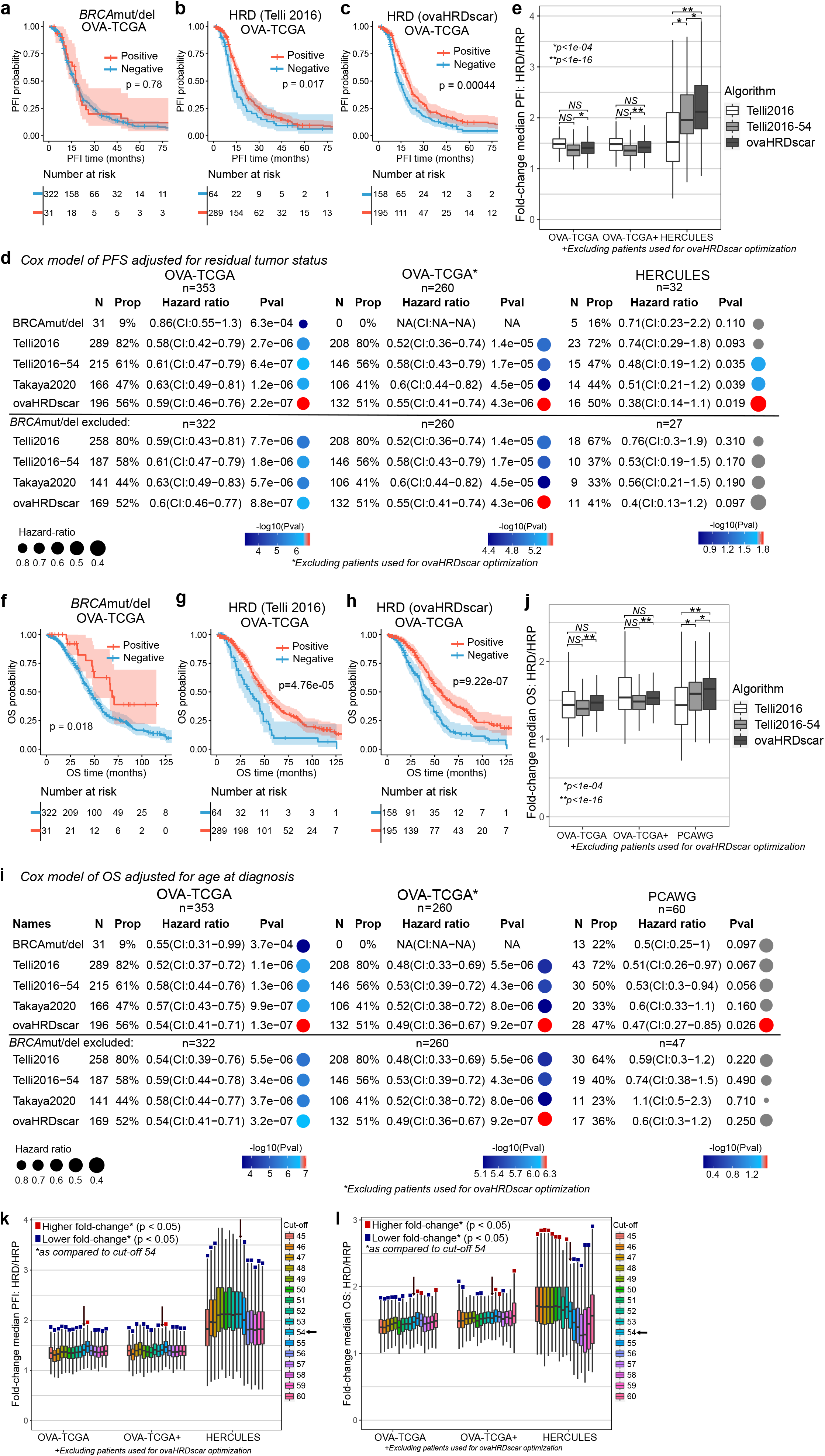
ovaHRDscar accurately predicts PFS and OS in HGSC patients. **a to c** Kaplan-Meier plots of PFS in OVA-TCGA patients stratified with different criteria, in **a** patients were stratified according to the *BRCA*mut/del status with no significant difference in their PFS probability over time (Log-rank, p=0.78); **b** patients were stratified according to the Telli2016 algorithm (Log-rank, p=0.017); and **c** patients were stratified using the ovaHRDscar algorithm. HRD patients had a prolonged PFS as compared to the HRP (Log-rank, p=4.4e-04). **e** Cox regression models for PFS adjusted for residual tumor after surgery according to the different HR classification criteria (*BRCA*mut/del, Telli2016, Telli2016-54, Takaya2020, ovaHRDscar). Three panels are shown: OVA-TCGA cohort in the left panel, OVA-TCGA cohort excluding the annotated HRD and HRP samples used for the detection of HRD-AIs in the middle panel, and the HERCULES prospective cohort (WGS) in the right panel. The number of patients (N) selected as HRD positive and their corresponding proportion (Prop), the hazard ratio for the Cox regression and the 95% confidence intervals (CI) and the p-value (Pval) of the regression are shown for each panel. The size of the dot represents the hazard ratio and color of the dot represents the p-value, grey dots represent non-statistical significant associations (p ≥ 0.05). **d** Fold-change of the difference in median PFS between HRD and HRP patients were stratified using ovaHRDscar, Telli2016 or Telli2016 using an HRD/HR cut-off value of 54 (Telli2016-54). Patients were bootstrapped 1000 times, and the fold-change was calculated for each iteration; the Box plots represent the values obtained by each bootstrapping iteration, no outliers are shown. U-test p-values are shown. **f to h,** Kaplan-Meier plots of OS for OVA-TCGA patients stratified using different criteria. **i** Fold-change of the difference in median OS between HRD and HRP patients stratified using ovaHRDscar, Telli2016 or Telli2016-54 using the same approach as in panel d. **j** Cox regression models for OS according to the HR-status classification criteria. The PCAWG samples in the right panel, the left and center panels are the same as in d.

Residual tumor after primary debulking surgery has been shown to be a strong independent prognostic factor in HGSC^20^. We next used residual tumor status as a covariable in Cox proportional hazard models to assess the performance of HRD algorithms in predicting the PFS. We found that ovaHRDscar positivity was significantly associated with prolonged PFS in OVA-TCGA also when adjusting for residual tumor (Wald test p=2.2e-07, **Fig. 3d**), similar to the Telli2016 (Wald test p=2.7e-06), Telli2016-54 (Wald test p=6.4e-07) and the Takaya algorithms (Wald test p=1.2e-06). The same was true also after excluding the annotated HRD/HRP samples used in the optimization (middle panel, **Fig. 3d**) and when not adjusting for the residual tumor (**Sup. Fig. 3d**). Importantly, ovaHRDscar significantly predicted PFS in the external HERCULES validation cohort (HR: 0.47 (CI:0.27-0.85), Wald test p=0.026). To compare how well the three algorithms (ovaHRDscar, Telli2016, Telli2016-54) can predict the differential outcomes of patients, we next calculated the differences in PFS between the HRD and HRP using a bootstrapping approach. Consistently, we found that the difference in PFS was significantly greater using the ovaHRDscar than using the Telli2016 algorithm in the independent HERCULES validation cohort (**Fig. 3e**). Moreover, ovaHRDscar was superior to the Telli2016-54 algorithm in the OVA-TCGA (**Fig. 3e**). In further validation, we inspected the performance of the HRD-classification algorithms in an additional independent prospective cohort (TERVA) with tumor-only SNP array profiling (see methods). Importantly, ovaHRDscar positivity significantly predicted longer PFS using Log-rank test and Cox proportional hazard model in the TERVA external validation dataset (**Sup. Fig. 3e to 3g)**.

We next explored the association of ovaHRDscar with overall survival (OS) in HGSC patients in the OVA-TCGA cohort and in an independent AU-OVA cohort in PCAWG (**Fig. 3f to h, Sup. Fig. 3h to j**). The clinical data in the prospective cohorts (HERCULES, TERVA) were not mature enough for OS evaluation. OvaHRDscar significantly predicted OS in the OVA-TCGA (**Fig. 3h**). In Cox regression analysis adjusted for age at diagnosis, ovaHRDscar significantly predicted OS, while the other algorithms did not reach statistical significance in the independent PCAWG validation dataset (**Fig. 3i**). These results were concordant also using a non-adjusted Cox regression analysis (**Sup. Fig. 3k**). Importantly, the median OS in patients with HRD tumors as compared to HRP was significantly longer when using the ovaHRDscar than using the Telli2016 or the Telli2016-54 algorithms in the independent PCAWG cohort when using a bootstrapping approach (**Fig. 3j**). Additionally, we compared the performance of ovaHRDscar to the CHORD algorithm that uses structural variation and a random forest implementation to classify HR-status^11^. In the PCAWG cohort, ovaHRDscar significantly predicted OS using the Log-rank test (**Sup. Fig. 3l, 3m**) and Cox proportional hazard models (**Sup. Fig. 3n),** while the CHORD algorithm did not show statistical significance.

Finally, to further investigate the impact of the ovaHRDscar cut-off value in predicting PFS and OS, we plotted the differences of median PFS and OS in HRD vs HRP when using different cut-off values in two independent validation test-sets (OVA-TCGA excluding samples used in the optimization and HERCULES) using bootstrapping (**Fig. 3k, 3l**). We observed that cut-off values lower than 54 led to significantly smaller differences (lower fold-changes) in PFS in the OVA-TCGA, and in the OVA-TCGA test-set, while higher values led to smaller differences in the HERCULES cohort (**Fig. 3k**). Further, values lower than 54 lead to smaller differences in

OS in the OVA-TCGA and OVA-TCGA test set, while higher values led to significantly smaller fold-change differences in the HERCULES cohort (**Fig. 3l**). Thus, the exploration of clinical outcomes in the multiple independent validation datasets supports HRD/HRP cut-off value of 54 as optimal for ovaHRDscar.

### Low intra-patient variation of ovaHRDscar in spatiotemporal tumor profiling

HGSC is characterized by high inter-tumor heterogeneity, and we next explored whether the anatomical site or timing of sample retrieval affects HR-status classification in HGSC. For this, we investigated the concordance of the ovaHRDscar levels in the HERCULES prospective cohort, which included 89 tumor samples from 33 HGSC collected from different anatomical sites and different treatment phases (treatment-naive, after neoadjuvant chemotherapy, or at relapse) (**Fig. 4a**). Consistent with the TCGA dataset, ovaHRDscar levels corresponded with the known genomic predictors of HRD (**Fig. 4b**). Importantly, we found that the levels were similar in paired, anatomically matched samples obtained before and after neoadjuvant chemotherapy, and also in primary (treatment-naive) versus relapsed tumors (**Fig. 4c**). Samples collected from different anatomical sites showed intra-patient variation (**Fig. 4a**), however it was lower than the observed inter-patient variation (U test p=1.95e-38**; Sup. Fig. 4a**). The intra-patient variability was not explained by differences in tumor purity (minimum 30%, see methods) (**Sup. Fig. 4b** and **Sup. Fig. 4c**). To determine the optimal anatomical sampling site, we next assessed HR-status per patient in treatment-naïve primary samples and compared ovaHRDscar calculated from different anatomical locations. Overall, the level of agreement for the HR-status classification ranged from 94% and 97% between the prioritization of different anatomical sites (**Sup. Fig. 4d**). However, ovaHRDscar status calculated primarily from ovarian or adnexal tumors was the strongest predictor for PFS (**Fig. 4d**, **Sup. Fig. 4e**). Consistently, prioritizing ovarian tumors accurately classified all tumors harboring *CCNE1* amplification as HRP in the prospective HERCULES cohort (**Sup. Fig. 4d**).

**Figure 4.**
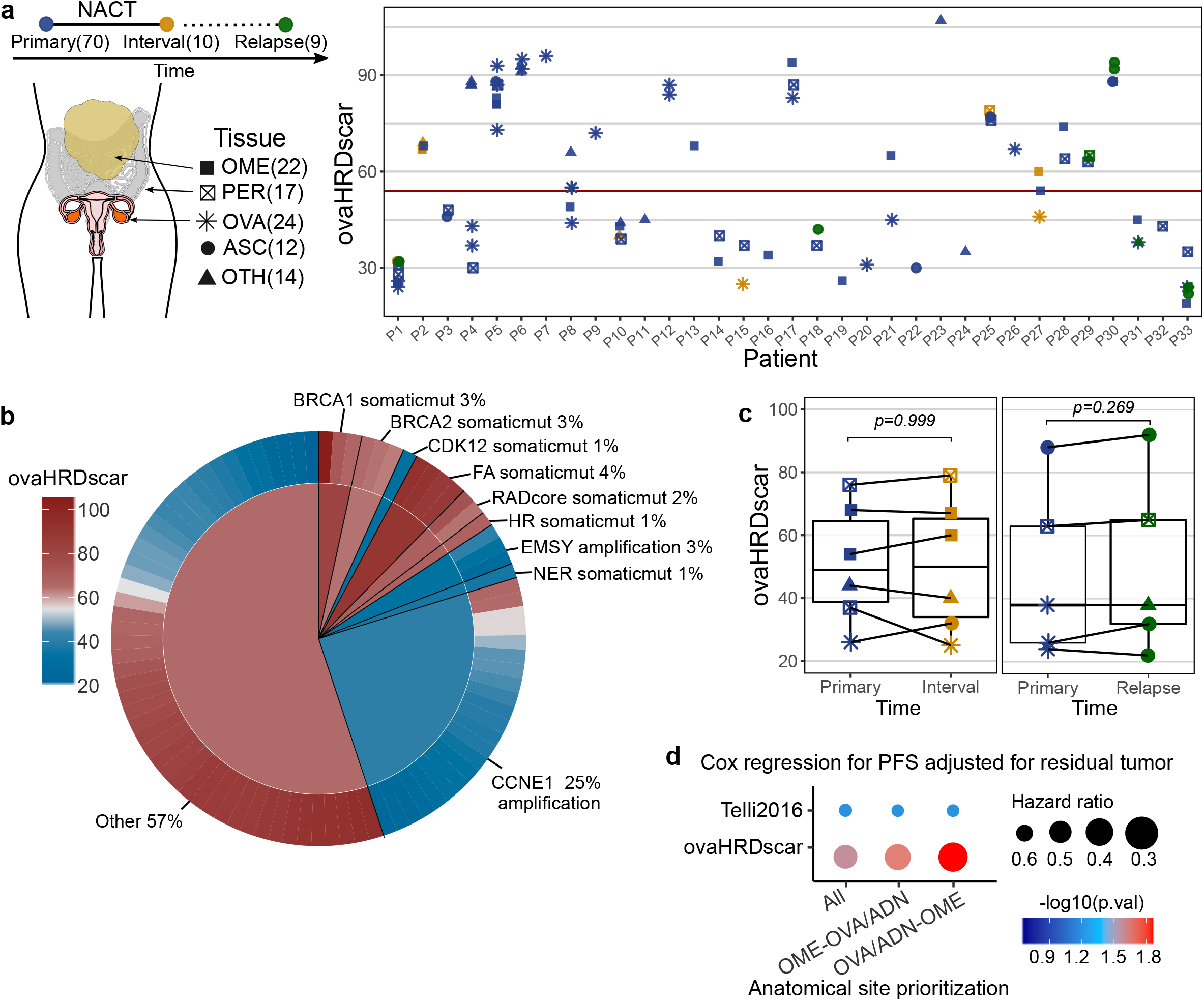
Intra-patient spatiotemporal variation of ovaHRDscar levels in 98 prospective HGSC samples. **a** Overview of the samples and their ovaHRDscar levels per patient in a prospective cohort (HERCULES). The tumor samples were collected at three different treatment phases and from different anatomical sites; the corresponding number of samples are displayed in parentheses. **b** Levels of ovaHRDscar in samples harboring different genetic or epigenetic alterations associated with HRD. The colors correspond to the ovaHRDscar levels, in the outer ring of the pie chart every bar represents a sample carrying the corresponding alteration, and average values for the genetic groups are displayed in the center of the pie chart. **c** ovaHRDscar values between paired samples for each patient (connected dots) did not change (Wilcoxon test) between the samples collected at different treatment phases. **d** Comparison of anatomical site prioritizations using Cox regression models for PFS using the Telli2016 or the ovaHRDscar algorithms. The size of the dot represents the HR and color of the dot represents the p-value. The HR-status for each patient is shown assessed using three anatomical sample prioritization approaches: 1) average HRD-AIs per all samples 2) omentum, and OVA/ADN if omentum sample not available (OME-OVA/ADN) 3) OVA/ADN, and then omentum if OVA/ADN not available (OVA/ADN-OME). In the case of multiple samples per same site, the average was used.

### Machine learning-aided detection of HRD-AIs improves the prediction of clinical outcomes in TNBC

Finally, we tested whether our systematic detection of HRD-AIs could improve previous algorithms when predicting clinical outcomes in TNBC. For this, using multi-omics data in TCGA and the same classification approach (**Fig. 2a**), we annotated 47 TNBC as HRD and 23 as HRP (**Fig. 5a**). Detection of HRD-LOH increased the accuracy of classification of HR-status from 80% (Telli2016 algorithm) to 93% (**Fig. 5b**). Likewise for LSTs, the accuracy increased from 93% to 98% (**Sup. Fig. 5a**) and for TAIs from 86% to 92% (**Sup. Fig. 5b**). Similarly as for the HGSC, instead of selecting TAIs of a particular length, we selected TAIs longer than 1Mb as this resulted in the largest increase in significance. The following HRD-AI criteria were observed as the most characteristic for TNBC: 1) LOH >10Mb and <30Mb, 2) for LSTs AI >5Mb with a distance between them <2Mb, and 3) TAI >1Mb. Then, using a subsampling approach, we identified that cut-off values for the sum of HRD-AIs (hereafter called tnbcHRDscar) from 47 to 53 produced the highest classification accuracy of the HRD and HRP samples (**Fig. 5c**), with the cut-off value of 53 as the closest value at the intersection of the HRP and HRP density distributions (**Fig. 5d**). Using the above criteria we observed that tnbcHRDscar increased the accuracy of classifying the HRD and HRP samples from 0.92 to 0.94 (**Fig. 5d**).

**Figure 5.**
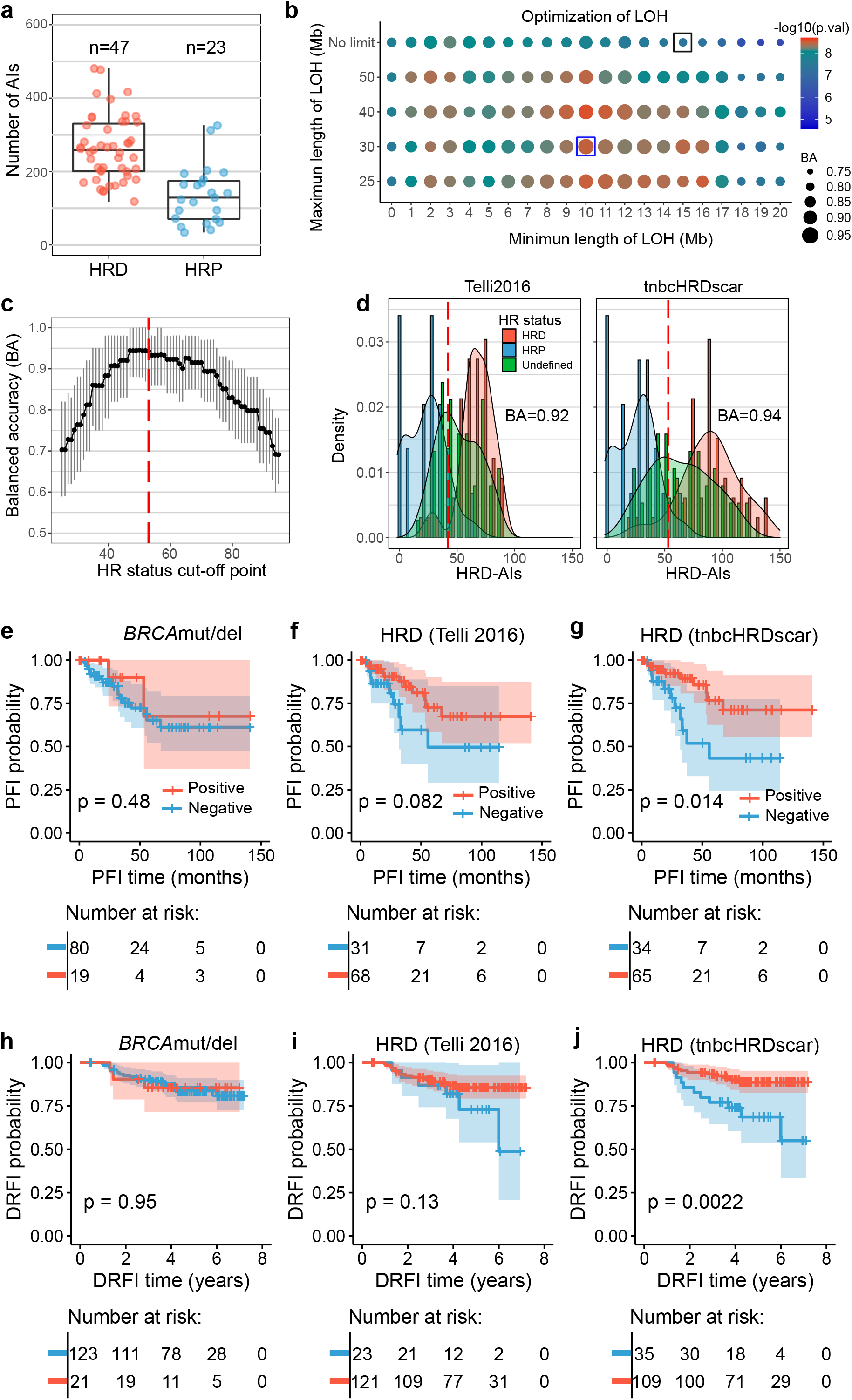
Machine learning-aided detection of HRD-AI in TNBC improves the prediction of clinical outcomes. **a** Number of AIs for TNBC in HRD and HRP samples in the TCGA. **b** Detection of LOH events. The size of the dots represents the decision tree balanced accuracy (BA) of classifying HRD and HRP using LOHs of the corresponding length, and the dot colors represent the difference in abundance of LOH between HRD versus HRP samples (U test, p value). Black box corresponds to the selection criteria utilized in the Telli2016 algorithm, and the blue box corresponds to the tnbcHRDscar BA and U test value. **c** Evaluation of the cut-off for tnbcHRDscar to define HR-status. The black dots connected with a line represent the balanced accuracy (BA) of the classification of the HRD and HRP samples using the given cut-off value, the 95% confidence intervals are shown in grey, the value of 53 (red dashed line) shows the highest BA. **d** Density distribution of HRD-AIs according to the Telli2016 and tnbcHRDscar algorithms. The red dashed line represents the cut-off established to define HR-status using Telli2016 (≥ 42) and tnbcHRDscar (≥ 53). The balanced accuracy (BA) for classifying the HR-status is shown for Telli2016 and ovaHRDscar algorithm. **e to g** Kaplan-Meier plots of PFS (Log-rank test) in TNBC patients in the TCGA stratified using: the *BRCA*mut/del status (**e**), the Telli2016 algorithm (**f**), the tnbcHRDscar (**g**). **h to j** Kaplan-Meier plots of distant relapse-free interval (DRFI, Log-rank test) of the TNBC patients in the validation dataset stratified using: the *BRCA*mut/del status (**h**), the Telli2016 algorithm (**i**), the tnbcHRDscar algorithm (**j**).

To test whether HR-status classification by tnbcHRDscar can predict clinical outcomes in TNBC, we next associated tnbcHRDscar with the PFS in the TCGA cohort and with the distant relapse-free interval (DRFI) in an independent TNBC SNP-array dataset^21^. Patients with the tnbcHRDscar-positive tumors had a significantly longer PFS than those with the tnbcHRDscar-negative tumors (Log-rank p=0.014), while *BRCA*mut/del status or the Telli2016 algorithm did not significantly associate with PFS (**Fig. 5e to 5g**). Only tnbcHRDscar showed a statistically significant association with the DRFI (Log-rank p=0.0022) in the independent validation dataset (**Fig. 5h to 5j**). Further, tnbcHRDscar classification in TCGA samples was also associated with OS (Log-rank p=0.039), similarly to the Telli2016 algorithm (Log-rank p=0.039; **Sup. Fig. 5c to 5e**). We next applied Cox regression analysis to validate the association of tnbcHRDscar with PFS and OS. In the TCGA cohort, tnbcHRDscar significantly predicted PFS (HR: 0.34, p=0.018, **Sup. Fig. 5f**) but the Telli2016 algorithm did not, while both similarly predicted OS (**Sup. Fig. 5g**). However, tnbcHRDscar but not the Telli2016 algorithm significantly predicted DRFI in the validation dataset (HR: 0.29, p=0.004, **Sup. Fig. 5h**). Additionally, we compared the performance of tnbcHRDscar with HRDetect^12^, an algorithm trained using WGS, to predict DRFI outcomes in the validation dataset. Interestingly, tnbcHRDscar improved the prediction of DRFI compared to the HRDetect (**Sup. Fig. 5h to 5j**), regardless of the cut-off values selected for the HRDetect (**Sup. Fig. 5k**).

## DISCUSSION

HRD tumors exhibit a distinct clinical phenotype with superior responses to platinum-based chemotherapy and sensitivity to PARP inhibitors. However, the accurate detection of HRD via somatic AIs has been confounded by the lack of systematic approaches and analyses performed in admixtures of tumor types with distinct genomic drivers. Herein, we established the HRDscar, a systematic approach for HRD detection to improve patient selection and clinical outcomes in cancer.

Several genomic approaches have been utilized to detect HRD, including 1) identification of single genetic mutations leading to predicted HRD^22^, 2) profiles of DNA repair deficiency gene expression^23,24^, 3) specific mutational patterns accumulated due to HRD^8,9,25^ or 4) structural genomic imbalances^7,26^. These genomic features have been implemented alone or in combinations in the search for optimal HRD detection, which has profound therapeutic implications^27^. It is now becoming accepted that benefits from the HR-directed therapies such as PARP inhibitors extend beyond the identification of HRD via individual genetic mutations^28^. This is due to the fact that genes such as *BRCA1/2* and *RAD51* paralogs can be altered beyond mutations via, e.g., hypermethylation or gene deletions^3,29^, and not all genomic events leading to HRD have yet been defined^30^. Allelic imbalances are indicative of the genetic consequences of HRD and, although not dynamically reflective of tumors’ functional HRD status, have shown promise as a biomarker predictive of the magnitude of benefit from PARP inhibitors, especially in the front-line setting^31,32^. The HRD-algorithm used in ovarian cancer clinical trials (Telli2016) was, however, generated using breast cancer samples or a mixture of breast cancer and ovarian cancer samples using *BRCA1/2* mutation as the sole determinant of HRD, and *BRCAwt* status as HRP^8,9,10^. Importantly, the European Society of Medical Oncology also indicated an urgent need to develop a more accurate HRD algorithm in HGSC to especially improve the identification of the HRP tumors^28^. Via a pan-cancer characterization of AIs, we discovered remarkable differences in the patterns of AIs of HGSC as compared to other cancer types, including TNBC, especially among the *BRCA*wt tumors. This prompted us to systematically identify the genomic footprints of HRD-AIs specific for HGSC using carefully annotated multi-omics data from TCGA and an iterative machine learning and statistical approach.

ovaHRDscar levels were concordant with tumor genetic alterations associated with HRD in the TCGA dataset and an external validation cohort (HERCULES). We found significantly lower levels of ovaHRDscar in tumors with *CCNE1* amplification, which was also previously proposed to be mutually exclusive with HRD and associated with poor clinical outcomes^33^. In line with a previous report^19^, tumors with *CDK12* mutation showed overall low levels of ovaHRDscar and thus could be considered HRP. In contrast, tumors with somatic mutations in *PTEN*, a gene associated with DNA repair^34,35^, showed high ovaHRDscar levels. However, the vulnerability of *PTEN* deficient cancers to PARP inhibitors remains to be verified in the clinical setting^28,36^. Further, ovaHRDscar showed a higher concordance with SBS3 than the Telli2016 algorithm. Most importantly, ovaHRDscar can be applied to detect HRD in HGSC samples using WGS or SNP-arrays, making it an attractive biomarker for the clinical setting.

A dichotomous thresholding of a predictive HRD biomarker is needed for therapeutic decision-making. In the Telli2016 algorithm, the cut-off for the total number of events was derived from a mixture of breast and ovarian cancer samples^10^. More recently, Takaya et al. set out to improve the HRD test by adjusting the cut-off value in ovarian cancer^19^. However, only *BRCA*mut status was used for separating HRD from HRP samples and the same genomic features of HRD-AIs were used as in Telli et al. In ovaHRDscar, after the development of accurate definitions of both the criteria of HRD-AIs and the cut-off, we identified more samples as being HRP, and separated HRD from HRP with improved accuracy over previous algorithms. When testing the Telli2016 algorithm using the ovaHRDscar cut-off value of 54, the accuracy was still below that of ovaHRDscar, indicating that both the accurate identification of the HRD-AIs and the selection of the optimal cut-off are needed to improve HRD detection in HGSC. In agreement, in most survival analyses, especially in the independent validation cohorts, ovaHRDscar outperformed the previous algorithms in predicting clinical outcomes.

HRD tumors are known to have superior responses to platinum-based chemotherapy and prolonged overall survival^37^. Consistently, ovaHRDscar improved the prediction of PFS and OS for platinum-based chemotherapy in the OVA-TCGA, also after excluding patients used when defining the criteria for ovaHRDscar. ovaHRDscar significantly predicted PFS and OS also among only the *BRCA*wt tumors. Importantly, ovaHRDscar improved the prediction of clinical outcomes in two independent patient cohorts and in multivariable models after adjusting for clinical covariables, indicating that ovaHRDscar reliably captures the phenotypic clinical behavior of HRD in HGSC. Further, using a disease-specific, systematic approach in the classification of HR-status, we could improve the prediction of the clinical outcomes also in TNBC, and tnbcHRDscar significantly predicted disease-free survival in the TCGA and in an independent dataset. However, none of the clinical cohorts included patients treated prospectively with, e.g., PARP inhibitors; therefore, prospective validation in larger patient series is warranted.

Finally, as HGSC is characterized by a high intra-tumor heterogeneity, we aimed at assessing whether the anatomical site of tumor sampling or the exposure to chemotherapy affects HRD detection. Our analysis of 98 samples collected from different anatomical sites and treatment phases indicated that ovaHRDscar levels remain similar within each patient, including anatomically site-matched samples collected before and after neoadjuvant chemotherapy. ovaHRDscar can thus be reliably assessed during routine clinical practice and also after neoadjuvant chemotherapy, given that the tumor purity remains higher than 30%. Interestingly, ovaHRDscar levels were also similar between treatment-naive and relapsed tumors, reflecting the nature of HRD-AIs as a historical consequence rather than a dynamic read-out of functional HRD. Analysis of different anatomical sites revealed that the overall inter-patient variation was higher than the intra-patient variation. However, in four out of 21 (19%) patients with samples from multiple anatomical sites, the HRD category depended on the anatomical site of sampling. The survival analyses indicated that ovarian or adnexal sites, followed by omentum, could be the preferred sites for HRD testing, warranting future validation in larger cohorts.

In conclusion, ovaHRDscar shows promise as a precise, clinically feasible assay for both outcome prediction and selection of patients for HR-directed therapies. With the fully documented, publicly available algorithms and generation pipeline, ovaHRDscar can be applied to other tumor types and implemented clinically for optimal patient selection to improve outcomes for patients with cancer.

## MATERIALS AND METHODS

### Data set collection and classification

For pan-cancer samples, allele-specific copy number segments were obtained from the Genomics Data Commons (GDC) portal (https://portal.gdc.cancer.gov/). The list of TNBC samples was adopted from Lehmann et al.^38^. For TNBC, samples were considered with *BRCA*mut if reported by Knijnenburg et al.^39^ to contain a gene deletion, gene mutation, or gene silencing of *BRCA*1 or *BRCA*2; while *BRCA*wt were considered those with no reported alterations.

For OVA-TCGA analysis, allele-specific copy number segments, DNA methylation, gene-level copy number profiles (including gene deletions), and clinical information data were obtained from the GDC data portal. Genes were considered with a “strong signal of deletion” if reported as such (labeled by −2) by Taylor et al.^40^. Gene promoter hypermethylation was considered when the probes up to 1500bp downstream of the transcription start site had an average beta value ≥0.75. The catalog of mc3 somatic mutations was obtained from the PanCanAtlas-GDC data portal (https://gdc.cancer.gov/about-data/publications/pancanatlas). Somatic mutations were classified according to the recommendations of the American College of Medical Genetics and Genomics^41^ using the web-tool VarSome^42^. Only pathogenic somatic mutations were considered in the analysis. For germline mutations, we selected those labeled as pathogenic and prioritized by Huang et al. 2018^43^. Genes were considered part of the HR pathway or other associated pathways according to the Kyoto Encyclopedia of Genes and Genomes database^44^. Complementary clinical information was obtained from the PanCanAtlas-GDC data portal. For PCAWG: allele-specific copy number segments, mutational drivers, and clinical information were obtained from the International Cancer Genome Consortium data portal (https://dcc.icgc.org/pcawg).

### Pan-cancer characterization of AIs

We used the allele-specific copy number segments from the Genomics Data Commons. Segments that did not span a whole chromosome and with a total copy number value different from two were selected as AIs. AIs shorter than 3Mb and longer than 50Mb were ignored. We quantified the number of AIs per sample and the median length of the AIs. The skewness of the distribution of the length AIs in different types of cancers was performed using the package DescTools.

### Selection of criteria for HRD-AIs

First, we annotated the OVA-TCGA samples as HRD and HRP according to the following. For HRD samples, samples harboring somatic or germline mutations, promoter hypermethylation, or strong signal of deletion of the genes *BRCA1, BRCA2*, and *RAD51* paralogs (**Fig. 2a**); for HRP sample annotation, we selected those with none of the HRD selection criteria, plus available data for methylation, gene deletion, somatic mutations and no deletion of any HR gene. The rest of the samples were annotated as “undefined” (**Fig. 2a**). The HRD and HRP annotation was used as “ground truth” in posterior accuracy assessment analysis. The HRP sample *TCGA-13-1511* was annotated as “undefined” as an outlier in the number of total AIs. Then, for the annotated HRD/HRP samples, we quantified the HRD-AIs (LOH, LST, TAI) according to Marquard et al.^45^ under different criteria. For LOH, we used length criteria (minimum length: *l_min_, maximum* length: *l_max_*). Exhaustively for each pair of values, *l_min_* and *l_max_*, we quantified the number of LOH per sample. We selected the pair of values that produced the highest classification power (see below) according to the HRD and HRP annotations. The quantification of LST events, defined by the parameters *s* (minimum AI length) and *m* (maximum distance between the AI events that comprise an LST event), was optimized similarly. Finally, we quantified TAI events if they were larger than *k*, where the length *k* was evaluated following the same approach. The classification power was evaluated by combining two approaches: 1) differential abundance of selected AIs in the annotated HRD vs HRP using one-tailed Mann-Whitney U test; 2) classification performance by decision trees (R package ‘rpart’) taking the abundance of the selected AIs as split-point. For the decision trees approach, samples above the split-point were tentatively considered as HRD and below - HRP, then true positive rate (TPR) and true negative rate (TNR) was computed when compared against the ground truth annotations (Fig. 2b). For each type of HRD-AI, we selected the set of parameters ({*l_min_, l_max_*}, {*s, m*}, *k*) with the highest product of U test p-value (p) and balance accuracy 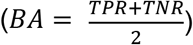 the product was inferred with the formula: −1 * log10(*p*) * *BA*. The selected set of parameters was incorporated in ovaHRDscar. The sum of HRD-AIs under the selected criteria was named the ovaHRDscar levels or values. A cut-off value to define the HR-status (samples with values above the cut-off are considered HRD and below - HRP) for ovaHRDscar and tnbcHRDscar levels was determined by exploring different cut-off values. For each cut-off value, we resampled with replacement 29 of the annotated HRD and 29 of the HRP cases 10,000 times; for each pseudo replicate, we calculated the balanced accuracy by comparing the HR-status using the cut-off value versus the ground truth annotations. Finally, we selected the cut-off value that produced the highest median balanced accuracy.

### Quantification of HRD-AIs

The quantification of HRD-AIs by the Telli2016 algorithm, the Takaya2020, the ovaHRDscar, and the tnbcHRDScar was performed using an in-house R-package (see code availability) adapted from the package scarHRD^46^. This package allows for the quantification of LOH, LSTs and TAIs under different selection criteria. Allelic imbalances smaller than 50bp were smoothed, as previously suggested by Popova et al.^7^. The selection criteria of HRD-AIs for Telli2016: LOH *l_min_* =15Mb, *l_max_ = 50Mb*; LSTs *s=12Mb, m=1Mb*, TAI *k*=1Mb, samples with HRD-AIs ≥ 42 were considered HRD otherwise - HRP. For the Takaya2020 algorithm, the same HRD-AIs selection criteria as for Telli2016 were used: samples with HRD-AIs ≥ 63 were considered HRD, and otherwise - HRP. For ovaHRDscar, the HRD-AIs selection criteria is: LOH *l_min_* =15Mb, *l_max_=50Mb;* LSTs *s=12Mb, m=1Mb*, TAI *k*=1Mb; samples with HRD-AIs ≥ 54 were considered HRD, and otherwise - HRP.

### Survival analysis

Survival plots, Log-rank and Cox regression models were performed in R using the packages “survminer” and “survival”. For OVA-TCGA, only patients disease treated with cisplatin or carboplatin were selected. For PCAWG, data from all patients were used (no treatment information available). Only data from primary samples (treatment-naive) were used. The *BRCA*mut/del status includes pathogenic somatic mutations, germline mutations, and “strong signal of deletion” in the genes *BRCA1* or *BRCA2*. Residual tumor after surgery was categorized as present or absent. For the indicated Cox regressions, residual tumor status or patient age at diagnosis was used as a covariable. Progression-free survival (PFS) and overall survival (OS) were defined as in Liu et al. 2018^47^. The CHORD signature HR-status classification for PCAWG samples was adopted from Nguyen et al. 2020^11^. In the TNBC cohort from TCGA, only patients with advanced Stage III-IV were selected. For survival analysis using HRDetect stratification, positive status was labeled for patients with an HRDetect value ≥ 0.7, and HRDetect negative for those with a value ≤ 0.2, patients with intermediate values were ignored. The mean differences of PFS and OS between HRD and HRP patients according to different criteria were calculated by bootstrapping the patients 1000 times; for each bootstrapping replicate was calculated the fold-change of median PFS or OS as median survival (PFS or OS) time in HRD patients divided by median survival (PFS or OS) time in HRP patients.

### Prospective HERCULES and TERVA data analysis

The tumor samples were prospectively collected in the HERCULES (http://www.project-hercules.eu) and TERVA (https://www.healthcampusturku.fi/innovation-new/terva-project/) projects. The Ethics Committee of the Hospital District of Southwest Finland approved both studies (Dnro: 145 /1801/2015). All patients gave their written informed consent to take part in the study. For HERCULES, paired fresh tumor and normal blood samples were sequenced using Illumina-HiSeq X Ten WGS. Raw reads were trimmed and filtered with Trimmomatic^48^, followed by duplicate marking with Picard Tools (https://broadinstitute.github.io/picard/). Alignment to the human genome GRCh38 was done using the Burrows-Wheeler aligner BWA-MEM^49^. Mutations were detected using GATK4-Mutect2 approach^50^. GATK4-Mutect2 was used for the detection of allele-specific copy numbers; regions listed in the ENCODE blacklist^51^ were omitted. Tumor purity was estimated using two approaches: 1) Based on somatic copy-number profiles using the software ASCAT v2.5.2^51^ 2) Based on variant allele frequency of the truncal mutation in gene *TP53* (*TP53*-VAF), purity was estimated using the formula: *2/((CN/ TP53-VAF) - (CN - 2))*, where CN corresponds to the absolute copy-number value estimated by ASCAT in the corresponding truncal mutation locus. Subsequently, the higher purity value was selected. For the TERVA samples tumor-only profiling, tumor samples were genotyped using the Infinium™ Global Screening Array-24 v2.0. B allele frequency and LogR ratios per sample probe were calculated using Illumina-GenomeStudio. ASCAT software was used for the detection of allele-specific copy numbers, *ascat.predictGermlineGenotypes* module was performed, adjusting parameters according to a panel of 200 normal germline blood samples. Intra- and inter-patient variability of ovaHRDscar values in the HERCULES cohort was determined by calculating the absolute value of the pairwise ovaHRDscar difference between all pair combinations of samples. Patient P19 was omitted from survival analysis because she received PARP inhibitors as maintenance after the first-line therapy.

### Statistics

The statistics analysis was performed in R. Difference in abundances was calculated using one-sided Mann-Whitney U test. Agreement was calculated using the Cohen kappa test. Concordance was measured using Lin’s concordance correlation coefficient. Level of correlations was assessed using Pearson correlations. P value less than 0.05 was considered statistically significant.

## Code Availability

The code used to detect HRD-AIs under different criteria is available on Github (https://github.com/farkkilab/findHRD-AIs). The ovaHRDscar algorithm implementation is available as an R package on Github (https://github.com/farkkilab/ovaHRDscar).

## Data availability

Data for the HERCULES and TERVA cohort will be available through the European Genome-Phenome Archive.

## ACKNOWLEDGMENTS

This study was funded by the Sigrid Jusélius Foundation (AF, LK), Cancer Society of Finland (AF, JC, LK), Academy of Finland (grant number 322979 to AF, grant numbers 314394 and 322178 to LK, grant number 314398 to SaHi), Paolo Foundation (AF), The Finnish Medical Foundation (AF), Finnish Cultural Foundation (AF), Instrumentarium Foundation (AF, JC), University of Helsinki (AF), AstraZeneca (to SaHi), the European Union’s Horizon 2020 research and innovation program under grant agreement No 667403 for HERCULES (SHa, JH, SaHi) and No 965193 for DECIDER (SHa, JH, SaHi). We also wish to thank FIMM Genomics core facility, The FINNPEC for the statistical reference data, Johan Staaf for assistance in TNBC data access, and the IT Center for Science (CSC) for computational resources.

## AUTHOR CONTRIBUTIONS

A.F. and F.P. conceived and designed the study with contributions from L.K., J.O., P.A.K. and S.H. F.P., J.O., J.C., D.C.G, Y.L. and K.L. analyzed the data. F.P. performed the statistical analysis. A.C., M.T., S.K., J. H., U. H., J.S.T. and H.L. performed experiments and provided materials. All authors wrote and approved the manuscript.

## COMPETING INTEREST

The authors declare no competing interests

Supplementary information is available for this paper

## SUPPLEMENTARY FIGURE LEGENDS

**Supplementary Figure 1.**
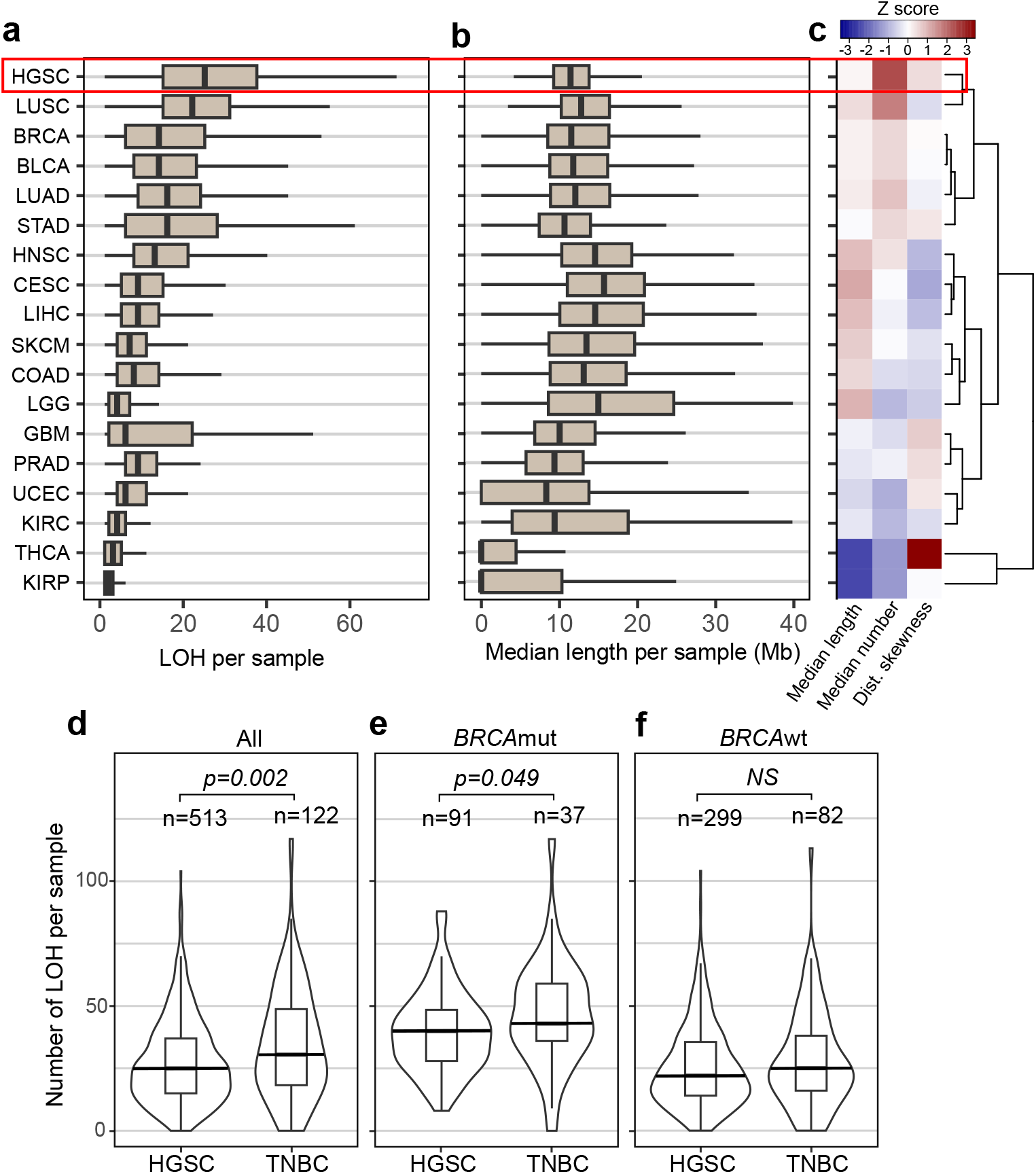
Pan-cancer characterization of LOH shows unique patterns in HGSC. **a.** Box plots showing the number of LOH events larger than 3Mb and smaller than 50Mb in the different cancer types. **b** Box plots showing the median length of LOH events (longer than 3Mb and smaller than 50Mb) in the cancer types. **c** Hierarchical clustering of the cancer types using the median length, median number of LOH events per sample, and the skewness of the distribution of LOH length. **d** Violin- and box plots representing the number of LOH events in all HGSC samples as compared to TNBC (U test, p=0.005). Long horizontal lines represent the medians. **e** Violin and box plots representing the number of LOH events in *BRCA*mut HGSC samples as compared to TNBC (U test, p=0.021). **f** Violin and box plots representing the number of LOH events in *BRCAwt* HGSC samples as compared to the TNBC (U test, NS).

**Supplementary Figure 2.**
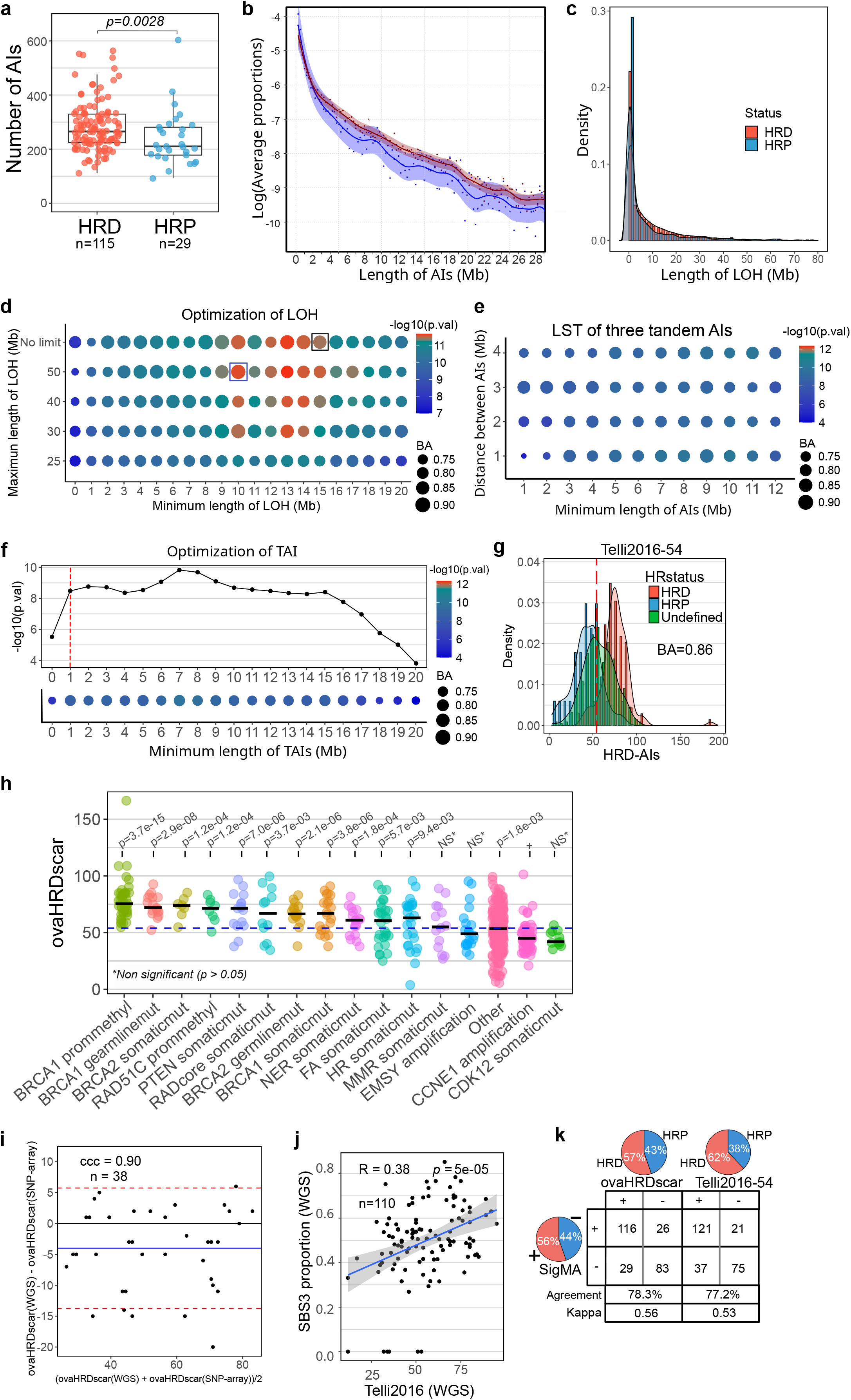
Descriptive statistics of HRD and HRP in HGSC. **a** Box plot showing the number of allelic imbalances in HRD (red) and HRP (blue) samples in OVA-TCGA. **b** The average proportion of segments (AIs) equal or greater than the given length in HRD (red) and HRP (blue) samples, the blue and red lines correspond to smoothing using cubic splines, confidence intervals are shown in shaded colors. **c** Density distribution of LOH events in HRD (red) and HRP (blue) samples. **d** The accuracy of the new LOH criteria (blue boxes) and those utilized in Telli et al. (black box); the size of dots represents the decision tree balanced accuracy (BA) when using the corresponding cut-off, colors correspond to the statistical difference in abundance of AIs between HRD versus HRP samples (U test p value). **e** Accuracy of HR classification using three tandem allelic imbalances LSTs of a given minimum length (x axis) and distance between them smaller than 1 to 4 Mb (y axis). Dot sizes and colors are presented similarly as in panel d. **f** Upper panel: Visualization of the statistical difference (U test p-values) in the abundance of TAIs between HRD versus HRP samples for selected TAIs length. Lower panel: The dot sizes and colors in the lower panel correspond to the description in panel d and e. **g** Density distribution of HRD-AIs according to Telli2016 algorithm, the red dashed line represents a cut-off vale of 54 to define the HR-status. The balanced accuracy (BA) of classifying the annotated HRD and HRP is shown, density distribution colors correspond to the samples annotated as in Fig. 2a. **h** OVA-TCGA samples stratified by genomic alterations and their corresponding ovaHRDscar levels. U test p values are shown for the comparison of ovaHRDscar levels between the corresponding alterations as compared to the samples with CCNE1 amplification. **i** Bland-Altman plot that shows the concordance (Concordance correlation coefficient, *CCC* = 0.90) between the number of ovaHRDscars detected using SNP-arrays and WGS in the intersecting samples from OVA-TCGA and PCAWG. **j** Correlation (Pearson, r=0.38) between the SBS3 proportion in WGS data from PCAWG versus the number of scars using the Telli2016 approach. Blue line shows the regression line, and the 95% confidence intervals are shown in grey. **k** The SBS3 status inferred using SigMA^16^ showing a higher agreement with ovaHRDscar (agreement=78.3%, Cohen’s kappa = 0.56) than with the Telli2016 algorithm using an HRP/HRP cut-off value of 54 (agreement=77.2%, Cohen’s kappa = 0.53). In the pie charts and table + and - correspond to the number of HRD positive and HRD negative samples identified under each criterion, respectively. On the bottom is shown the number of samples and the level of agreement between the corresponding criteria.a

**Supplementary Figure 3.**
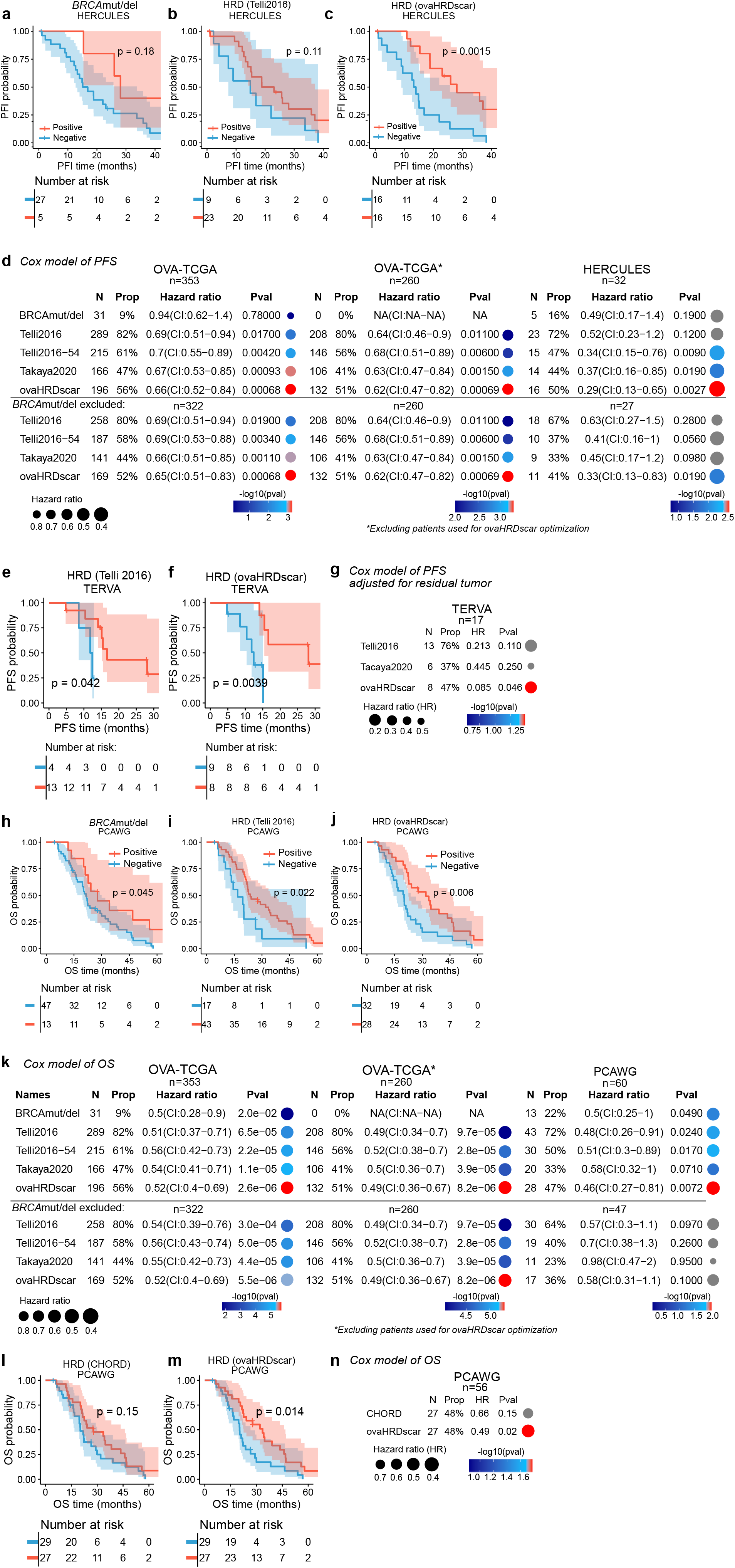
ovaHRDscar shows an improved prediction of PFS and OS in HGSC patients. **a to c** Kaplan-Meier plots for PFS in the HERCULES cohort. The patients were stratified using different criteria: The *BRCA*mut/del status in the left panel (**a**), The Telli2016 algorithm in the middle panel (**b**) and The ovaHRDscar algorithm (**c**). **d** Cox regression models for PFS in HGSC patients using different selection criteria. The colors, rows and columns descriptions are the same as in Figure 3d. **e to f** Kaplan-Meier plots of PFS in the prospective TERVA cohort. The patients were stratified using: The Telli2016 algorithm on the left (**e**), the ovaHRDscar on the right (**f**). **g** Cox regression models for PFS in the TERVA cohort. The patients were stratified using different criteria. The colors, rows and columns descriptions are the same as in Figure 3d. **h to j** Kaplan-Meier plots for OS in the HERCULES cohort stratified using the different criteria: *BRCA*mut/del status on the left (**h**), the Telli2016 algorithm (**i**) and the ovaHRDscar algorithm (**j**). **k** Cox regression models for OS adjusted for patient age at diagnosis in HGSC patients stratified using different criteria similarly as in Figure 3h. **l to m** Kaplan-Meier plots for OS in the PCAWG cohort stratified using: the CHORD signature on the left (**l**), the ovaHRDscar on the right (**m**). **n** Cox regression models for OS in the PCAWG cohort. Patients were stratified using CHORD and ovaHRDscar criteria; the dot colors, rows and columns descriptions are the same as in Figure 3d.

**Supplementary Figure 4.**
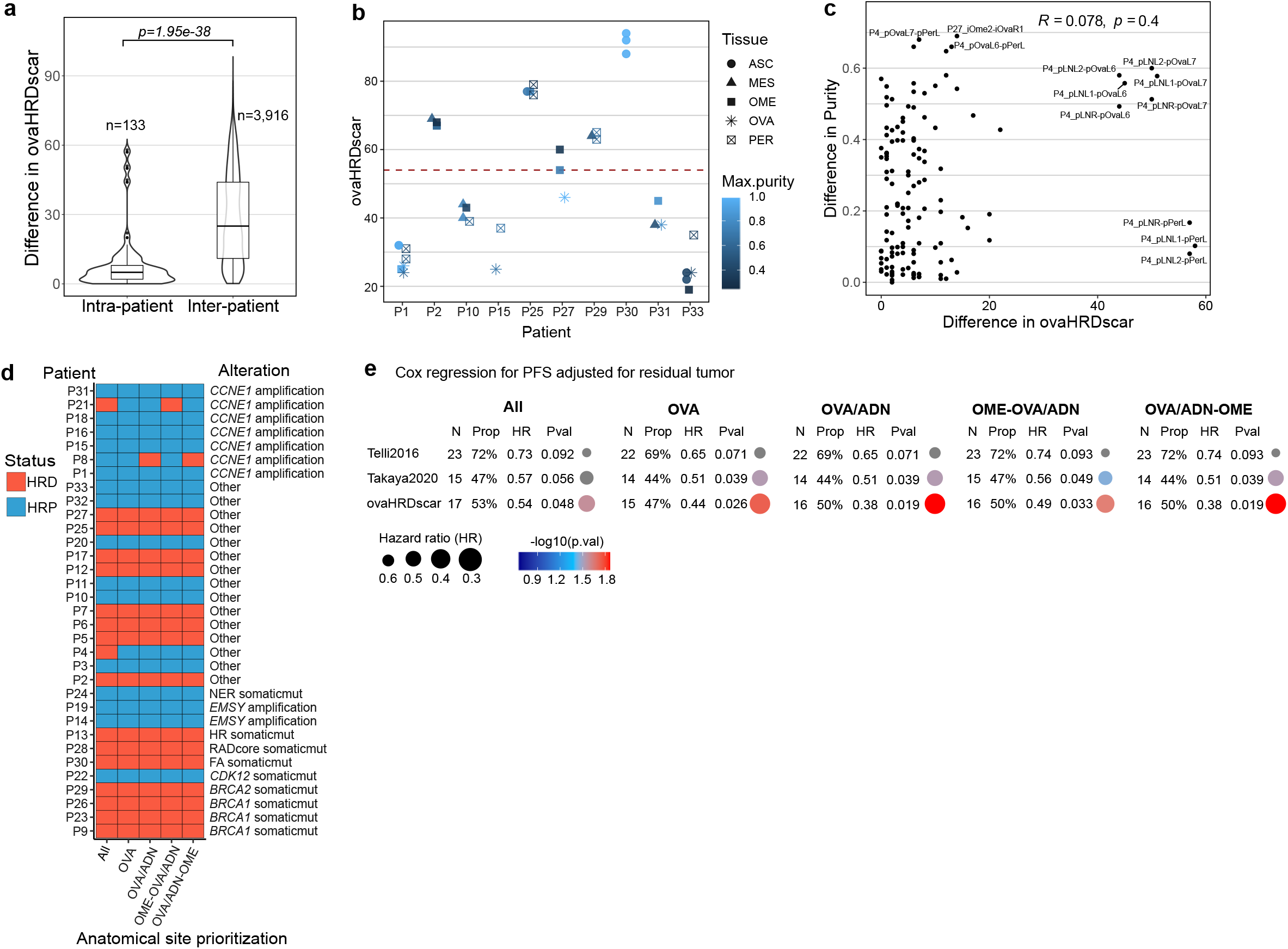
Inter and intra-patient variability of ovaHRDscar levels. **A** Left box plot for the difference of ovaHRDscar values between all possible intra-patients’ samples pairs. Right box plot for the corresponding difference of ovaHRDscar values in all possible inter-patients’ samples pairs in the HERCULES prospective cohort. **b** Samples from different anatomical sites with tumor purity and ovaHRDscar levels indicated. **c** The correlation between the difference of ovaHRDscar values between all intra-patient sample pairs and the difference in tumor purity of the corresponding sample pairs. Patient P4 was ignored as an outlier. **d** Color table of HR-status classification and HR-related genomic alterations using five different approaches to prioritize anatomical sites for ovaHRDscar calculations. **e** Cox regression models for PFS adjusted for the residual tumor after surgery in the HERCULES cohort using different algorithms and five different anatomical site prioritizations. Colors, rows and column descriptions are the same as in Figure 3d.

**Supplementary Figure 5.**
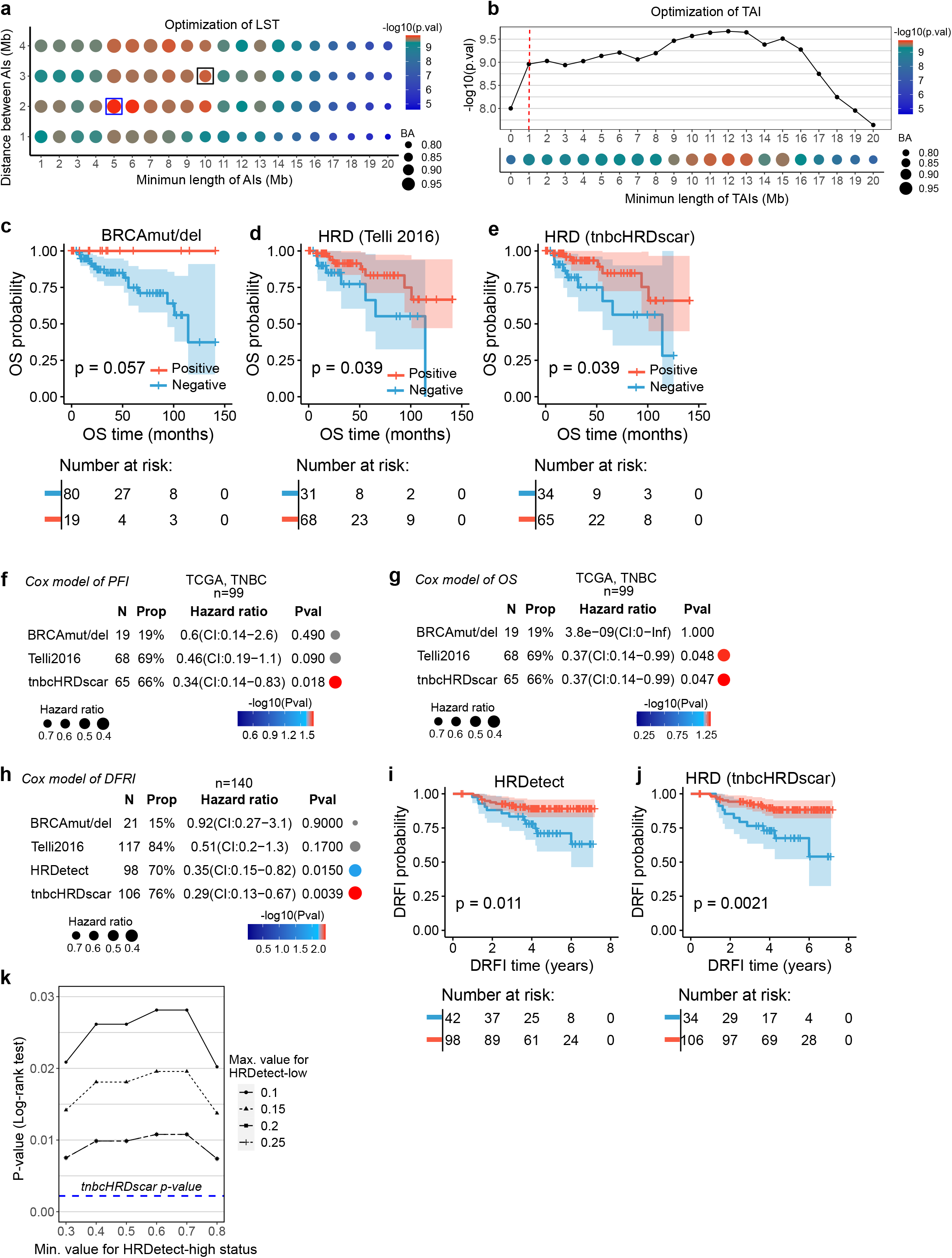
Machine learning-aided detection of HRD-AI in TNBC improves the prediction of clinical outcomes. **a** Generation of HRD-LST events. The size of the dots represents the decision tree balanced accuracy (BA) of the classification of HRD and HRP when selected LSTs with the corresponding criteria. The dot colors correspond to the statistical difference in abundance of the selected LSTs between HRD versus HRP samples (U test, p-value). The black box corresponds to the selection criteria proposed by Telli2016, blue box corresponds to the tnbcHRDscar BA and U test value. **b** Upper panel: Visualization of the change in p-values (U test) when selecting TAIs >1Mb (red dashed line). Lower panel: the difference in abundance of TAIs of selected length between HRD versus HRP samples. The dot sizes and colors in the lower panel correspond to the description in panel a. **c to e** Kaplan-Meier plots for OS in the TCGA’s TNBC patients stratified using the different criteria; **c** *BRCA*mut/del status on the left, **d** the Telli2016 algorithm, and **e** the tnbcHRDscar algorithm. **f** Cox regression models for PFS in HGSC patients stratified using different criteria, the dot colors descriptions are the same as in Figure 3d. **g** Cox regression models for OS in in the TCGA’s TNBC patients stratified using the different criteria. **h** Cox regression models for DRFI in an independent TNBC patient cohort^21^ stratified using different criteria. **i to j** Kaplan-Meier plots for DRFI in an independent TNBC patient cohort^21^ stratified using: HRDetect based on whole genome sequencing data (**i**), the tnbcHRDscar based on SNP-array data (**j**). **k** Selection of different values to define the HRDetect-high/low status for patient stratification in the TNBC patient cohort^21^ and its association with DRFI (Log-rank test, p-value). Patients with intermediate HRDetect values were ignored. In blue line, the Log-rank p-value when using tnbcHRDscar in panel **j**.

